# High-throughput screening reveals mechanisms of environmental control of germination in a fungal thermophile

**DOI:** 10.64898/2025.12.04.692316

**Authors:** Olusola A. Ogunyewo, Kristopher Fleming, Madeleine Morris, Kayleigh Fort, Lori B. Huberman, Rachel B. Brem

## Abstract

*Thermothelomyces thermophilus* is a filamentous fungus isolated from self-heating compost. Unlike most of the fungal kingdom, this species exhibits a growth optimum at 45°C and is intolerant of temperatures below 30°C. To investigate genetic contributors to temperature-dependent fitness in this system, we implemented a large-scale insertional mutagenesis approach. We generated thousands of *T. thermophilus* mutants and cultured them at temperature extremes in standard medium. Phenotyping-by-sequencing identified dozens of disrupted loci representing candidate determinants of thermophilic life history, including several annotated in metal transport. We then validated a subset of screen hits with a directed, single-gene knockout paradigm. The results revealed a temperature-dependent regulatory logic for germination, the developmental decision by which a fungal spore initiates growth. Surprisingly, most mutants germinated far better at 50°C than the wild-type in standard medium and showed markedly slower germination at lower temperatures, consistent with altered germination regulation rather than enhanced intrinsic heat tolerance. We hypothesized that *T. thermophilus* has evolved sophisticated regulatory machinery to block germination at high temperature unless environmental conditions are favorable. As a proof of concept, we surveyed media conditions and established that elevated zinc dampened germination of wild-type *T. thermophilus* at 50°C but promoted it at lower temperature; mutation experiments made clear that such sensitivity was mediated in part by the zinc transporter *zip*. We interpret these results under a model in which *T. thermophilus* integrates temperature and nutrient availability to control the transition from spore dormancy to vegetative growth, a developmental decision that shapes fitness outcomes across temperatures.

**Significance:** Fungal thermophiles thrive at temperatures that represent the upper limits of eukaryotic life. The regulatory and developmental mechanisms that shape their temperature-dependent fitness remain poorly understood. In this work, we elucidate how *Thermothelomyces thermophilus* integrates temperature cues with other environmental inputs during germination, a key life-cycle stage for dispersal. Our findings highlight germination regulation as an important contributor to fitness at elevated temperatures in a thermophilic eukaryote. These insights are of basic biological interest and provide a foundation for rational strategies to modulate temperature-dependent performance in industrial strains, with applications for high-temperature bioprocessing.

## Introduction

Thermophilic fungi operate at the upper temperature range of eukaryotic life, with a physiological ceiling approaching ∼60°C (1). Microbial ecology surveys have outlined the narrow diversity of truly thermophilic fungal species adapted to composts and geothermal soils (2, 3). In thermophilic bacteria, elevated growth temperatures have been linked to genome streamlining, higher GC content, thermostable protein folds, membrane lipid saturation, and enhanced chaperone and DNA-repair capacities (4, 5). Whether similar mechanisms govern thermophily in fungi remains unclear, particularly given the added complexity of organelles, chromatin regulation, and developmental programs in these systems. Thus, against the backdrop of years of work on mesophile and thermotolerant fungi (6–9), the mechanisms of high-temperature growth preference in fungal thermophiles are still incompletely understood.

Beyond their fundamental biological interest, thermophilic fungi also have practical relevance as potential hosts for industrial bioprocessing. Microbial cell factories are widely engineered to produce fuels, pharmaceuticals, and other industrially valuable chemicals; however, the cost of cultivation substrates remains a major constraint to sustainable biomanufacturing (10, 11). Fungi represent a potential solution toward this end, when they have the capacity to digest abundant, low-cost crop wastes like rice straw and corn stover. For such substrates, industrial processes commonly apply high-temperature pretreatments to accelerate biomass deconstruction. In this pipeline, subsequent cooling prior to microbial inoculation adds cost and energy demand (12). Thermophilic fungi therefore represent a promising chassis for bioproduction, as they could bridge pretreatment and fermentation steps without the need for cooling, directly converting pretreated biomass into target products.

Among thermophilic fungi, *Thermothelomyces thermophilus* (formerly *Myceliophthora thermophila*) has emerged as a compelling model with robust growth near 50°C and a carbohydrate active enzyme repertoire capable of plant cell-wall deconstruction, sparking interest in its use as a thermophilic cell factory (13, 14). Genetic tools in this system now enable precise engineering (15), including the potential to optimize high-temperature performance(16–18). We set out to use *T. thermophilus* as a testbed for the dissection of mechanisms of fungal thermophily at a genomic scale. Our approach was to pioneer the application of a recently developed method for randomly barcoded insertional mutagenesis with screening via sequencing in filamentous fungi (19) for the *T. thermophilus* system. We sought to use this strategy for an unbiased genetic survey of temperature-responsive growth behaviors in *T. thermophilus*. We used the results as a jumping-off point for exploration of the temperature dependence of the asexual stages of the filamentous fungal life cycle and the underlying genetics. The results reveal how a thermophilic eukaryote coordinates stress signaling and development at high temperature.

## Results

### Temperature-dependent growth and development of *T. thermophilus*

With the ultimate goal of understanding temperature-dependent growth and developmental behaviors in *T. thermophilus*, we first surveyed growth behaviors of *T. thermophilus* across temperatures and conditions. On solid agar, we documented the asexual growth cycle, from hyphal expansion through formation of aerial hyphae to production of spores (conidia) with visible pigmentation (Figure 1A). We observed a temperature-responsive lag in development as temperature increased, even with high-percentage agar and high-humidity incubation to maintain plate integrity (Figure 1A-B). As a complement, we also assayed growth in liquid culture, in which we inoculated *T. thermophilus* conidia into standard liquid medium and quantified the resulting mycelial biomass accumulation after incubation with shaking. The results revealed evidence for *bona fide* thermophilicity by *T. thermophilus*, with an optimum at 45°C during liquid growth and a gradual falloff of performance under warmer and cooler conditions (Figure 1C). We thus considered liquid culture as a growth format suitable for genetic analyses to reveal mechanisms contributing to temperature-dependent fitness in *T. thermophilus*.

**Figure 1.**
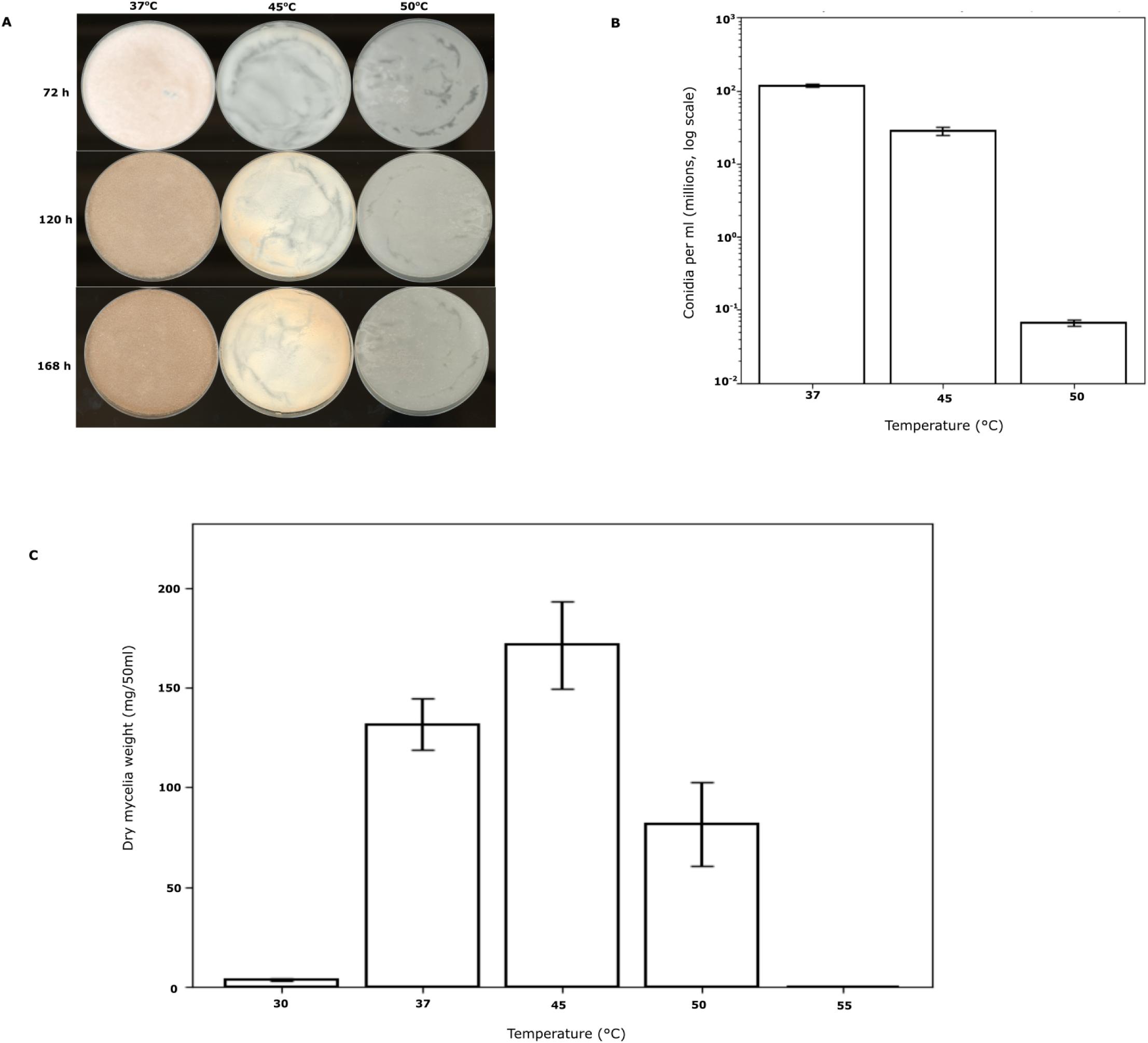
Growth and developmental measurements of *T. thermophilus* across temperatures. (A) Colony morphology of the wild-type strain grown on solid Vogel’s medium at 37°C, 45°C, and 50°C and imaged at 72 h, 120 h, and 168 h post-inoculation. (B) Conidial density measured from cultures grown at 37°C, 45°C, and 50°C after 168 h. The *x*-axis indicates growth temperature (°C), and the y-axis indicates conidia per milliliter (log scale). Bars represent mean ± SD from three biological replicates. (C) Dry mycelial biomass measured from 50 ml liquid cultures incubated for 24 h at the indicated temperatures. The *x*-axis indicates incubation temperature (°C), and the *y*-axis indicates dry mycelial weight (mg per 50 ml culture). Bars represent mean ± SD from three biological replicates.

### Construction and mapping of a *T. thermophilus* insertional mutant library

To develop a genomic resource for mapping of genotype to phenotype in *T. thermophilus*, we constructed a pool of barcoded insertional mutants. Our process used random insertion from *Agrobacterium tumefaciens* harboring hygromycin-marked, barcoded transfer DNA (T-DNA) (19), with which we carried out ectopic transformation of mononucleate *T. thermophilus* conidia via established procedures (20) with minor modifications to conidial density, co-cultivation duration, and selection conditions, followed by expansion and aliquoting of replicate sub-pools (Figure 2A). To catalog the diversity and chromosomal distribution of T-DNA insertions across the mutants, we isolated DNA from replicate cultures of the pool grown in liquid medium at 37°C and prepared sequencing libraries enriched for T-DNA–genome junctions. *In silico* analysis of reads mapped to the *T. thermophilus* reference genome identified the T-DNA right-arm junction, extracted adjacent genomic sequence, and assigned uniquely mapped barcodes to genomic coordinates. From these data, we detected 17,155 distinct insertion sites across all seven chromosomes. Their distribution was largely uniform with a few prominent peaks (Figure 2B); most notable of the latter was a T-DNA insertion hotspot on chromosome 7 corresponding to an rRNA-encoding locus (Supplementary File 2), suggestive of high transcriptional activity as a driver of insertion prevalence. In total, 3,426 T-DNA insertions were located within annotated genes (Supplementary File 1), establishing our pool as a partial-coverage mutant collection suitable for quantitative fitness profiling in *T. thermophilus*.

**Figure 2.**
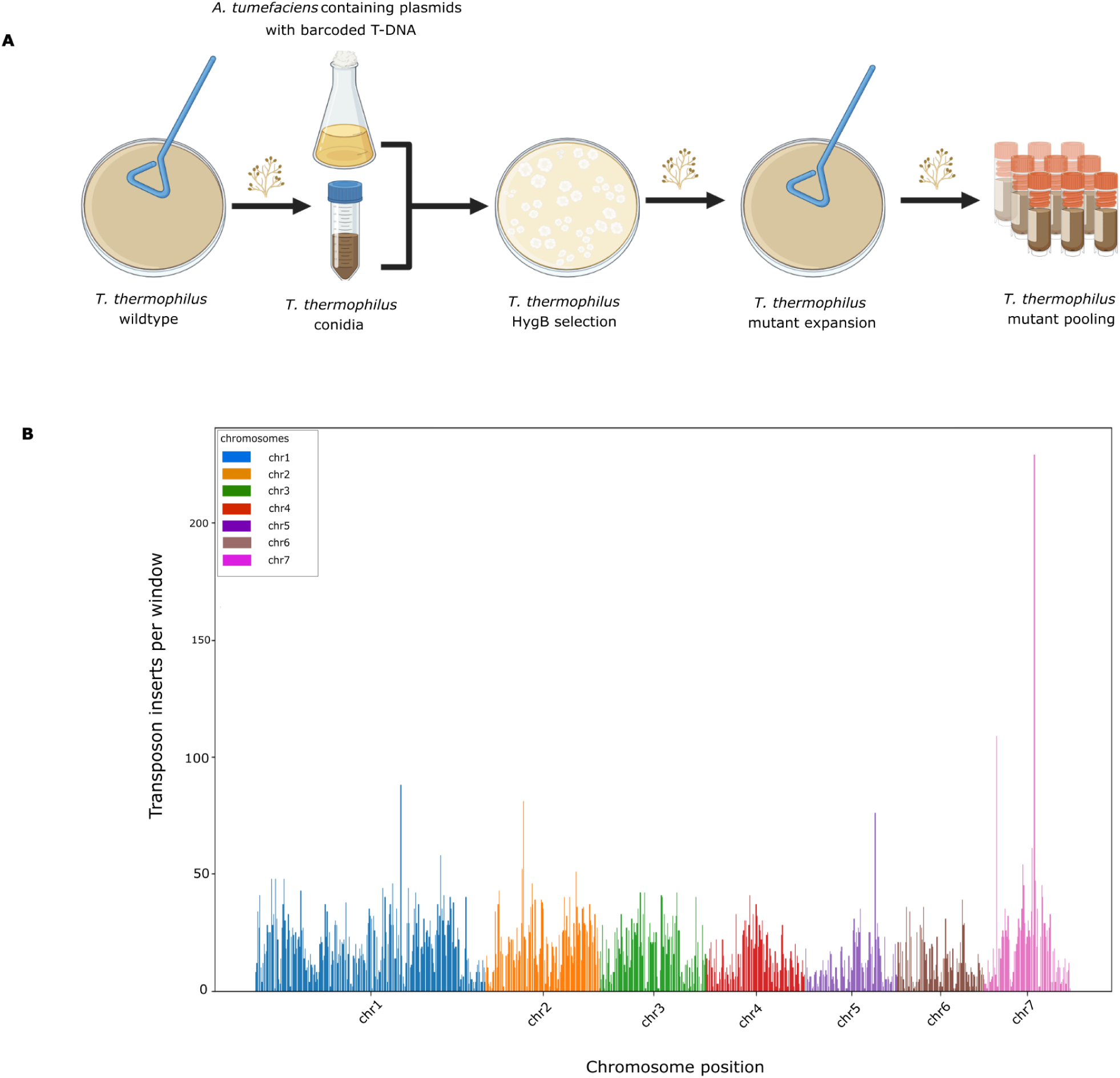
Construction and genomic distribution of the *T. thermophilus* barcoded insertional mutant library. (A) Overview of the Agrobacterium-mediated transformation workflow used to generate the barcoded insertion mutant library. *T. thermophilus* conidia were co-cultivated with *A. tumefaciens* containing plasmids with barcoded T-DNA. Transformants were selected on hygromycin-containing medium. Transformant colonies were bulked by allowing them to conidiate on selection plates, and conidia from all plates were harvested and pooled to generate the barcoded insertion mutant library (B) Genome-wide mapping of T-DNA insertions across the seven chromosomes of *T. thermophilus*. The *x*-axis indicates chromosomal position (grouped by chromosome), and the *y*-axis indicates the number of mapped T-DNA insertions per 50 kb genomic window. Colors correspond to individual chromosomes.

### Genome-wide quantification of temperature-dependent mutant fitness

We expected that screens with our insertional mutant pool could enable dissection of temperature-dependent fitness in *T. thermophilus*. To this end, we sought to use the resource to identify genes whose mutation effect on fitness depended on temperature. We designed an experimental scheme using growth assays of our mutant pool at 37°C and 50°C, representing temperatures that approached the lower and upper extremes, respectively, of wild-type temperature tolerance in liquid culture (Figure 1C). For each biological replicate, we cultured independent aliquots of the *T. thermophilus* mutant pool in parallel at 37°C and 50°C, and we quantified mutant abundance by sequencing the barcode regions of the inserted T-DNA from the resulting genomic DNA (Figure 3A). In the resulting abundance data, reproducibility between biological replicates was high across both temperatures (Pearson r > 0.994; Figure 3B and Supplementary Figure S1). We used the complete data set as input into a quality control and analysis pipeline (see Methods) that assessed differences between temperatures in mutant abundance on a per-gene basis, finding 783 mutants at adjusted *p*-value < 0.05 and absolute log_2_(fold-change) ≥ 1 (Supplementary File 3). Though these genes spanned a range of annotated functions (Supplementary File 3), their behavior in the screen was remarkably consistent: for all but six gene hits, mutant abundance was higher at 50°C than at 37°C (Figure 3C and Supplementary File 3). These data attested to a highly complex genetic architecture of temperature response in *T. thermophilus,* whose contributors, when mutated, almost invariably conferred defects in cooler conditions and/or benefits at high temperature.

**Figure 3.**
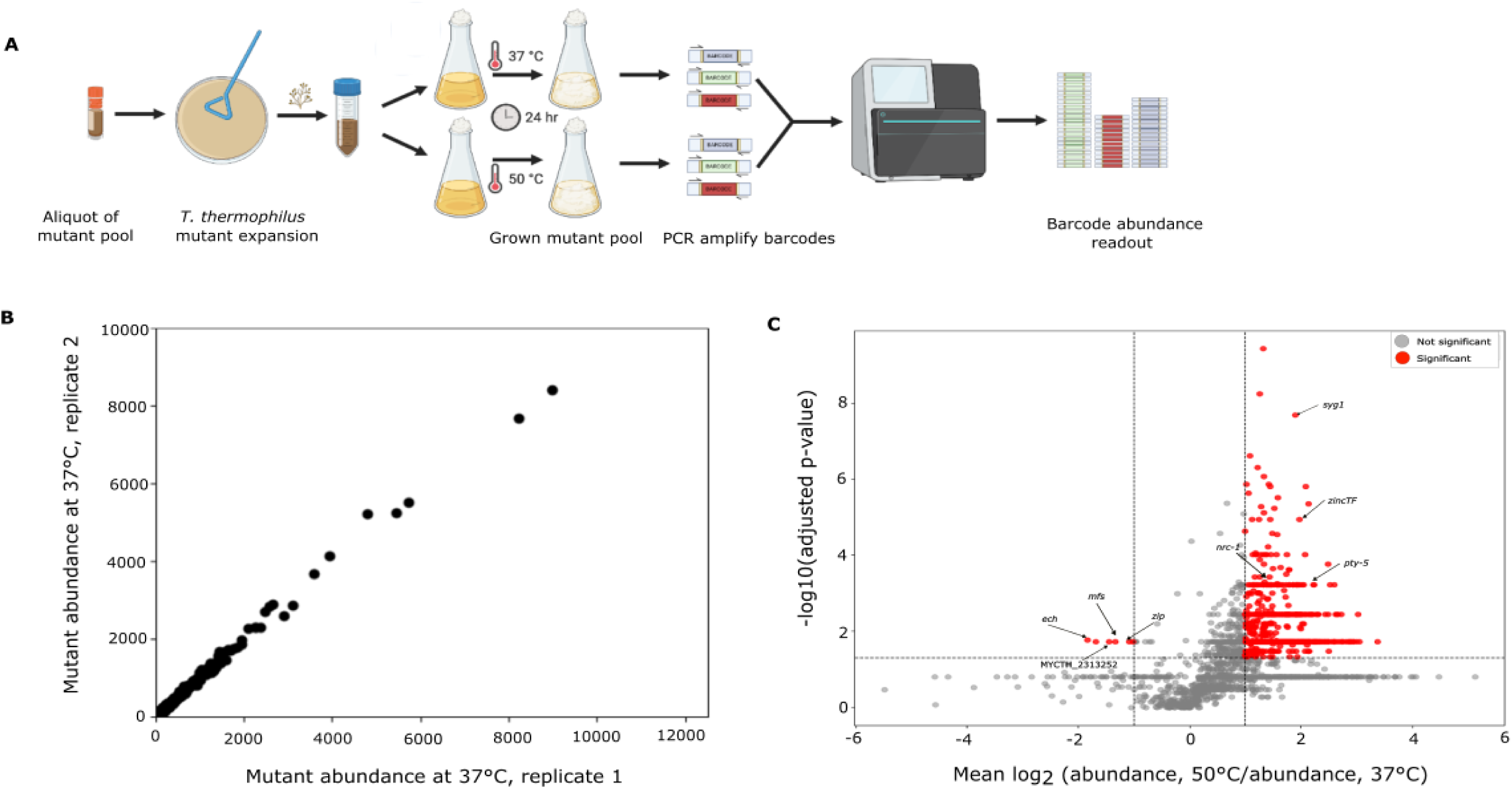
Genome-wide BarSeq fitness profiling identifies temperature-dependent mutant defects. (A) Experimental workflow for barcode sequencing (BarSeq) of the *T. thermophilus* mutant pool. Conidia were inoculated from the pool stock onto solid medium, harvested, and used to start liquid cultures. Cultures were incubated for 24 h at 37°C or 50°C. Genomic DNA was extracted from each culture, and barcode regions were amplified and sequenced. Sequencing reads were used to quantify barcode abundance for downstream fitness analysis. (B) Example comparison of mutant abundances between two biological replicates grown at 37°C. Each point represents one mutant, and the *x*- and *y*-axes report normalized abundance in the indicated replicate. (C) Each point reports the results of a test for differential abundance of insertional mutants in one gene between temperatures. The *x*-axis reports the log of the ratio of two quantities: the average abundance, across genotypes and biological replicates, of mutants at the respective gene at 50°C, and the analogous quantity at 37°C. The *y*-axis indicates – log₁₀(adjusted *p*-value) from statistical testing. Points in red indicate loci meeting quality control thresholds for temperature-dependent abundance change in terms of significance, effect size, and variability (see Methods). Genes chosen for single-gene follow-up are highlighted and labeled with arrows.

### Single-gene validation of temperature-dependent screen hits

To validate the inferences from our genome-wide random mutagenesis screen, we took a single-gene approach. For this purpose we chose eight screen hits (Figure 3C and Table 1) with high coverage in the mutant pool and robust mutant effects favoring either 50°C or 37°C. These included putative regulators (*zincTF, syg1*, *pty-5*, and *nrc-1*), transporters (*zip* and *mfs*), and housekeeping genes (*ech* and *MYCTH_2313252*) (Table 1). We generated targeted deletion mutants for each by homologous recombination of a hygromycin-resistance cassette at the endogenous *T. thermophilus* locus in a strain harboring a deletion in *ku80*, to ensure efficient deletion by eliminating non-homologous end joining. Correct integration events were confirmed by PCR and sequencing (Supplementary Figure S2).

**Table 1.** Hits from high-throughput screen subjected to single-gene validation. Each row reports effects of mutation of one gene on temperature-dependent fitness as measured by large-scale pooled T-DNA mutagenesis, from Figure 3C, for a locus chosen for follow-up single-gene validation analysis. For each gene, the table reports the *T. thermophilus* gene identifier (GeneID); the corresponding gene symbol; the annotated protein name; the log of the ratio of two quantities: the average abundance, across genotypes and biological replicates, of mutants at the respective gene at 50°C, and the analogous quantity at 37°C; and the adjusted *p*-value from a Wilcoxon rank-sum test for differential abundance of the respective mutants between temperatures.

| GeneID | Gene symbol | Protein name | Mutant effect on abundance, $\log_2(50^\circ\text{C}/37^\circ\text{C})$ | Adjusted *P*-Value |
| --- | --- | --- | --- | --- |
| MYCTH_2139072 | *zincTF* | Zn(2)-C6 fungal-type domain-containing protein | 1.98 | 1.12 E-05 |
| MYCTH_2300403 | *pty-5* | Tyrosine-specific protein phosphatase-domain-containing protein | 1.95 | 5.84 E-02 |
| MYCTH_100809 | *syg1* | SPX domain-containing protein | 1.90 | 2.01E-08 |
| MYCTH_2305107 | *nrc-1* | MAPKK kinase | 1.15 | 8.6 E-04 |
| MYCTH_116881 | *mfs* | MFS general substrate transporter | -0.98 | 0.0184 |
| MYCTH_2303755 | *ech* | Enoyl-CoA hydratase | -1.40 | 0.0078 |
| MYCTH_2307966 | *zip* | Zinc/iron permease | -1.45 | 0.0184 |
| MYCTH_2313252 | MYCTH_2313252 | Putative UPF0658 Golgi apparatus membrane protein | -1.83 | 0.0171 |

In designing experiments to characterize these single-gene mutant strains, we reasoned that temperature could influence multiple stages of the *T. thermophilus* life cycle, and that distinguishing between such effects would maximize our insights into the mechanisms of temperature response. We elected to focus on the phases relevant for the growth scheme of the high-throughput screen (Figure 3), namely germination of conidia and the extension of hyphae after vegetative growth had established. To this end, we established an assay to quantify temperature impacts on conidial germination of a given strain by microscopy, at a relatively early timepoint where germlings could easily be resolved from one another in an imaging field (Figure 4A). Separately, we designed an assay of temperature-dependent vegetative hyphal extension for a given strain, avoiding confounding effects from germination by using a standardized pre-germinated inoculum (Figure 4B). We applied both assays to each mutant and to their isogenic parent, the otherwise wild-type Δ*ku*80 strain. This scheme allowed us to distinguish effects on germination from those on hyphal growth once a colony was established, in the interpretation of mutant phenotypes.

**Figure 4.**
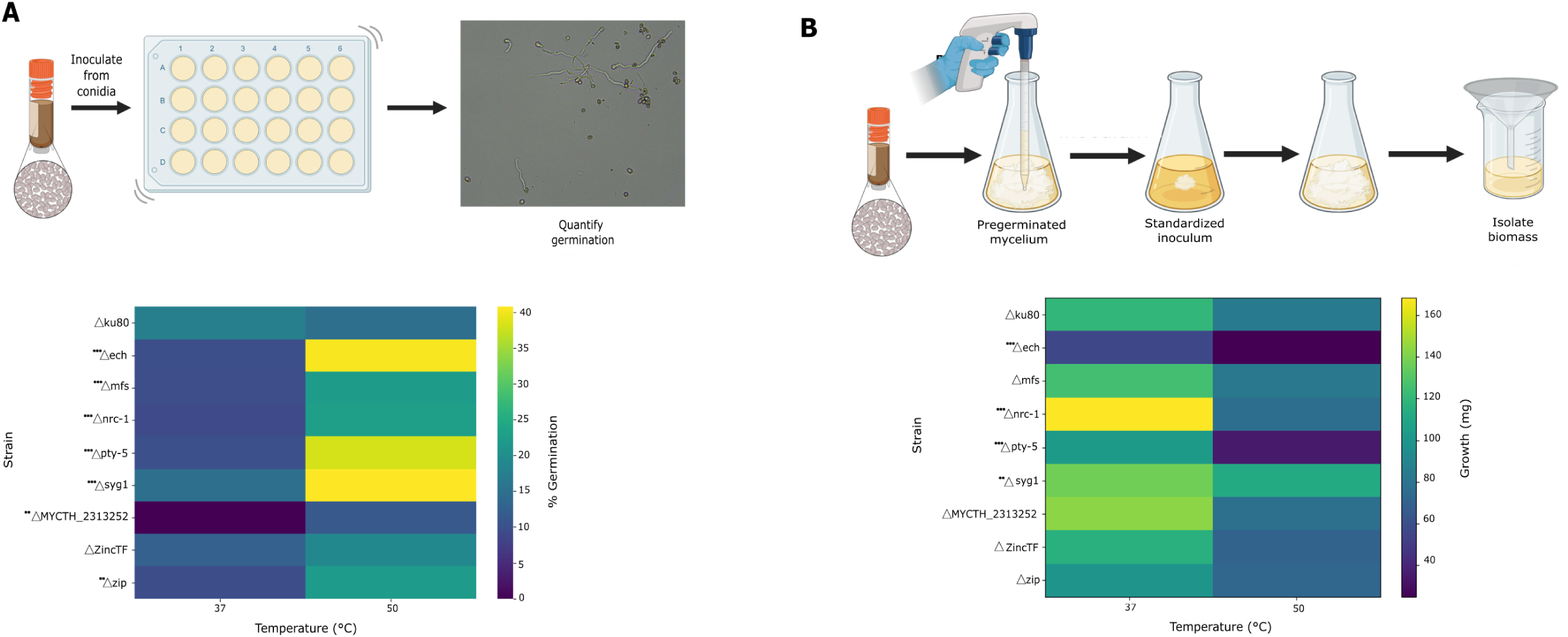
Germination and biomass measurements for temperature-tested single-gene deletion strains. (A) Top, germination assay. Conidia were inoculated into microtiter wells and incubated at a temperature of interest, followed by scoring via microscopy. Bottom, each row reports results from germination assays of the indicated *T. thermophilus* strain at the indicated temperatures. Each cell reports the average, across replicates, of the percent of conidia scored as germinating. (B) Top, biomass assay. Conidia were inoculated into liquid medium and incubated to allow germination and pre-growth; the resulting mycelium was used as inoculum into a second culture, followed by incubation at a temperature of interest, and biomass harvest and quantification. Bottom, each row reports results from growth assays of the indicated *T. thermophilus* strain at the indicated temperatures. Each cell reports the average, across replicates, of dry biomass (mg) for each strain at each temperature. Statistical significance was evaluated using a two-way ANOVA (strain × temperature) with post-hoc comparisons relative to the Δku80 reference strain at each temperature. Asterisks denote significance (\**p* < 0.05; \*\**p* < 0.01; \*\*\**p* < 0.001).

From germination measurements across this strain set, we observed a modest drop in performance at elevated temperature by the Δ*ku80* parent strain, which we used as a baseline for comparison of the remainder of the single-gene mutants (Figure 4A). Among these strains, 6 trends mirrored results from our large-scale screen, in which fitness phenotypes of mutants made by random integration almost universally favored high abundance at 50°C (Figure 3B). Interestingly, in three exceptions to the latter (*mfs*, *ech,* and *zip*), for which mutant abundance in the screen was highest at 37°C (Figure 3B and Table 1), our single-gene validation experiments nonetheless established that the respective mutants germinated preferentially at 50°C. This observation, and the fact that a single-gene deletion of *MYCTH_2313252* conferred a germination defect at the lower temperature (Figure 4A) but random insertional mutants at this locus in the screen were highly abundant at 37°C relative to 50°C (Figure 3B and Table 1), suggested cases in which performance in the growth scheme of the screen reflected contributions from behaviors other than those interrogated in our germination assay. However, apart from Δ*MYCTH_2313252*, the most salient and pervasive pattern from our observations of germination in single-gene mutants was that of improvements at high temperature (Figure 4A). We thus formulated a model in which the wild-type alleles of many loci function as repressors of germination at high temperature, such that their deletion would release this brake and allow development to proceed despite heat stress; the same loci would act as activators of germination under cooler conditions.

We next turned to assays of biomass accumulation in the hyphal growth phase of *T. thermophilus*, inoculating with pre-germinated mycelium at an equal starting concentration for all strains, including the *Δku80* parent (see Methods). For wild-type *T. thermophilus*, high temperature retarded growth in this scenario, with 1.5-fold lower biomass accumulation at 50°C than at 37°C (Figure 4B). Most single-gene mutants phenocopied this behavior almost exactly, such that we identified few cases in which a mutant grew differently from the wild-type parent at either temperature (Figure 4B). Among these exceptions, we noted an advantage of the *nrc-1* deletion mutant relative to wild-type at 37°C; defects at 50°C in the Δ*pty-5* and Δ*ech* strains (representing a plausible mechanism for increased abundance of mutants at these loci at 37°C in our original large-scale screen [Figure 3B and Table 1]); and increased growth by Δ*syg-1* at 50°C (Figure 4B). But as the latter represented outliers across the gene set, we concluded that growth was not a primary life cycle stage for the roles of most of our focal genes in temperature response, particularly in light of the striking impact of their deletions during germination (Figure 4A). Given the origin of our focal genes as representative of the hundreds of hits from our large-scale screen (Figure 3B), we infer that the overall trends in our screen results were also likely dominated by temperature-dependent effects of gene disruption on germination rather than vegetative growth.

### Zinc availability modulates germination efficiency

Is there a regulatory logic that governs repression of germination in wild-type *T. thermophilus* at high temperature? We hypothesized that the fungus integrates multiple environmental inputs, including temperature and nutrient availability, in the decision to germinate. As a test case, we considered environmental metal exposure, in light of the validated role in temperature-dependent germination of the putative zinc-iron permease *zip* (Figure 4A). We formulated a model in which zinc availability acts to gate *T. thermophilus* germination under heat stress. To explore this, we performed a zinc titration series for wild-type *T. thermophilus*, assaying germination as a function of zinc concentration and temperature. The results revealed a dose-responsive benefit in germination from increasing zinc at 37°C and the inverse relationship at 50°C, with germination falling off markedly as zinc concentrations increased (Figure 5A). At the lowest zinc concentrations, germination at high temperature far outstripped that observed at 37°C (Figure 5A). Such a pattern dovetailed with the benefit to germination at 50°C that we had observed in standard medium (5 mg/L zinc) upon deletion of *zip* (Figure 4A), as expected if zinc uptake were limited in the Δ*zip* mutant. To test the role of *zip* directly, we repeated germination assays in the Δ*zip* strain and found no impact of zinc concentration on germination efficiency in this background (Figure 5B). Rather, germination remained higher at 50°C than at 37°C across zinc conditions in the absence of *zip* (Figure 5B). Together, these data demonstrate that zinc is a key environmental input into developmental decision-making in *T. thermophilus*, and that zinc-dependent modulation of germination requires *zip*.

**Figure 5:**
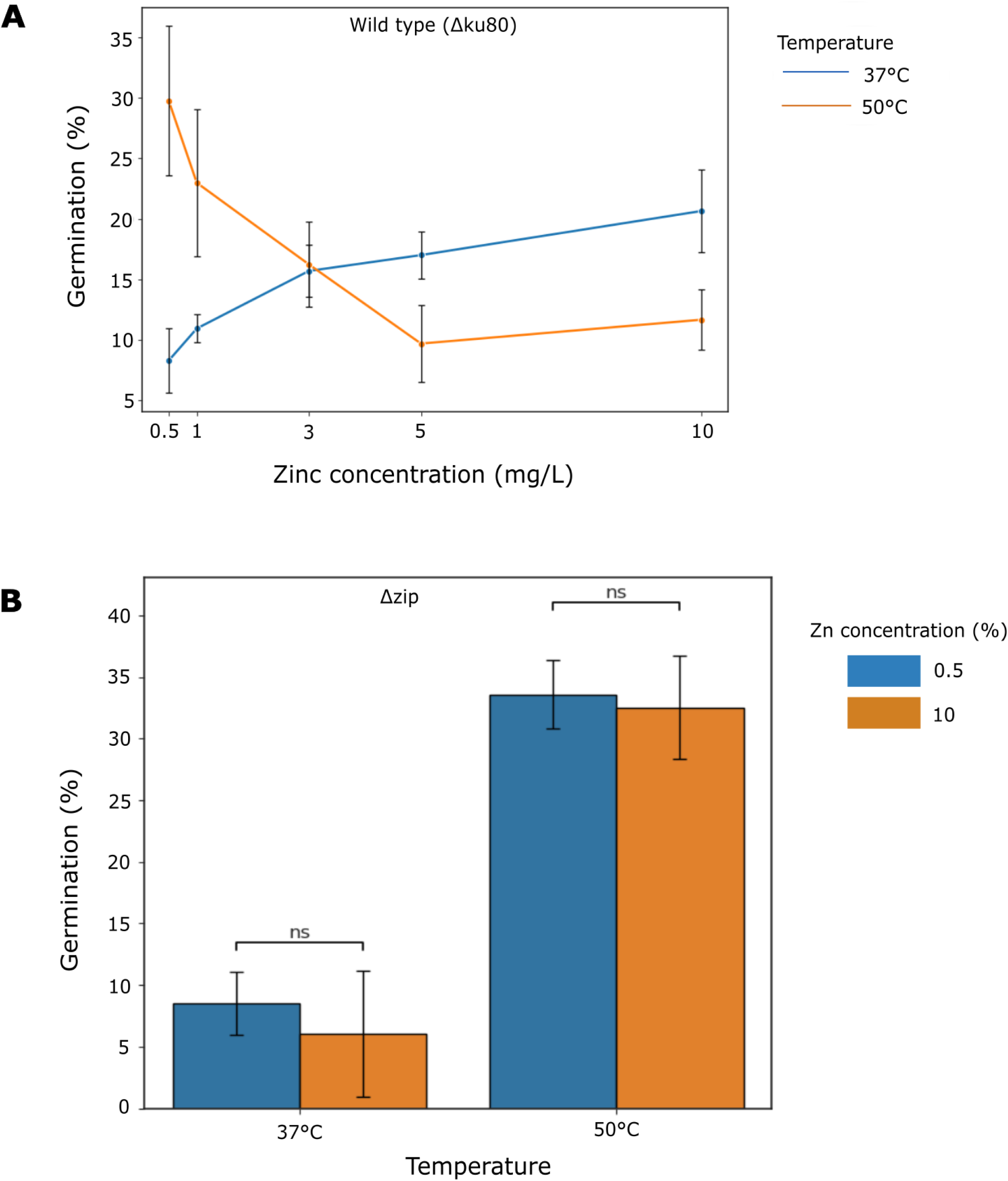
Zinc-dependent modulation of germination in *T. thermophilus*. (A) Germination of wild-type *T. thermophilus* across zinc concentrations and temperatures. The *x*-axis indicates zinc concentration in the medium (mg/L), and the *y*-axis indicates the percentage of germinated conidia. Each point represents the mean value, and error bars denote standard deviation across biological replicates. (B) Germination of *T. thermophilus* Δ*zip* across zinc concentrations and temperatures. The *x*-axis indicates temperature and the *y*-axis is as in (A). Each bar represents the mean value, and error bars denote standard deviation across biological replicates. ns, not significant in a *t*-test comparing germination between zinc concentrations for the indicated temperatures.

## Discussion

Thermophilic fungi are a powerful model for the basic biology of how eukaryotes adapt to extreme environments, and they also are of interest in industrial applications. In this work, we have established insertional mutagenesis as a tool for mechanistic dissection of temperature response in *T. thermophilus.* High-throughput forward genetics in filamentous fungi is in its infancy (19), and our method is well-positioned to accelerate advances in the *T. thermophilus* system, as has now become standard in bacteria and yeasts (21–25). The principles that have emerged from our study break new ground in the understanding of the thermophilic life history for fungi.

We have shown that hundreds of genes, when disrupted in our high-throughput screening approach, affect high-temperature performance by *T. thermophilus.* Among these, our validation of focal gene hits has found that most act as negative regulators of conidial germination at high temperature, in that they improve germination when deleted. The capacity to make spores represents a mechanism for stress resistance common to essentially all fungi, and exit from this dormant state is tightly regulated even in mesophiles (26–28), to ensure that a new colony only commits to growth under favorable conditions. It is tempting to speculate that for thermophiles, evolution has optimized the dormancy-breaking decision in unique ways, taking into account the cost or toxicity of growth programs at extreme temperatures. If so, the logic in *T. thermophilus* would be to sense environmental signals associated with the worst risk at high temperature and use them to trigger a block on germination.

Our screen and validation data raise the possibility that zinc processing represents one such environmental risk factor. If cellular functions involving zinc were particularly deleterious during *T. thermophilus* germination under high-heat conditions, it would mirror previous studies in mesophilic fungi in which metal exposure compromises spore germination more than it does hyphal growth (29). Plausibly, such effects could be mediated by reactive oxygen species, which have been associated with both zinc (30, 31) and elevated temperature (32, 33) in other systems. Formally, however, our data leave open the question of whether and how zinc exerts damaging effects *per se* in *T. thermophilus*. What our experiments have shown is that levels of zinc that shut down germination at high temperature fostered the best germination at 37°C. This pattern is consistent with a regulated delay of germination under combined zinc and heat stress, rather than solely a passive consequence of zinc-induced damage; indeed, the zinc concentrations we have tested here fall within ranges commonly used in fungal physiology studies (41–43). That said, we see zinc as only one example of an environmental cue that can be integrated with thermal signals to shape developmental decisions by *T. thermophilus*. We expect that the annotated regulator genes whose deletion boosts germination at high temperature will ultimate prove to represent nodes of a complex signal-integration network controlling the *T. thermophilus* life cycle. Such a function could reflect repurposing from the known roles, in mesophiles, of our focal genes *pty-5* (a tyrosine phosphatase) and *nrc-1* (a MAP3 kinase) in conidiation and germling fusion respectively (34, 35); the function of another of our focal factors, *syg-1,* likely in phosphate-based signaling (36), is unknown even in model fungi. For the annotated effector genes we have highlighted here—*ech* (encoding enoyl-CoA hydratase, a β-oxidation enzyme whose disruption alters growth and development in other fungi) and the zinc/iron permease gene *zip* (37, 38)—we interpret their mutant effects as eliminating repressive environmental inputs farther upstream, allowing germination to proceed even under heat stress.

Looking ahead, although we have emphasized relief of germination repression in *T. thermophilus* mutants, our work has also uncovered genes for which we infer positive effects on mycelial growth (focal genes *ech* and *pty-5*): that is, their deletion compromises this phase of the life cycle at high temperature. We anticipate that many more such pro-growth factors remain to be identified in *T. thermophilus*. It is reasonable to assume that principles from thermophilic bacteria and archaea that lack multiple life stages, in which evolution has adjusted housekeeping genes all over the genome to promote growth at high temperature (44), are likely to bear out in fungi. Our screening approach for *T. thermophilus* may have been particularly likely to miss pro-growth loci of this kind if they are essential, in that their disruption mutants would fail to propagate in our stocks at any temperature and would be lost from our analysis pipeline. As such, and in light of the partial coverage of our mutant pool, our screen results represent only a first window onto the complete suite of genes that mediate the thermophilic life history of *T. thermophilus*. As they stand, however, our findings underscore the importance of germination as a nexus of regulatory decisions for *T. thermophilus*, integrating temperature with other environmental inputs—which could well include nutrient availability, pH, or osmotic or oxidative stress, as well as metal homeostasis—to optimize fitness.

## Conclusion

We have used an insertional mutagenesis screen and validation follow-up in *Thermothelomyces thermophilus* to uncover principles of thermophilicity in fungi. Our results shed light for the first time on the regulatory landscape of decisions by these organisms about whether to germinate in the heat. And in an industrial setting, the deletion mutants we study here represent compelling candidate cases in which genetic manipulations could improve performance by *T. thermophilus* even at the extreme of its heat tolerance.

## Methods

### Strains, media, and growth conditions

*Thermothelomyces thermophilus* ATCC 42464 was obtained from the American Type Culture Collection. The Δ*ku80* strain MJK20.2 was generously provided by Vera Meyer (45, 46). The strains were maintained on Vogel’s minimal medium (VMM) (47) supplemented with 2% sucrose. For solid cultures, VMM with the indicated carbon source was solidified with 3% (w/v) agar. Liquid cultures were grown in 250 mL Erlenmeyer flasks at 200 rpm with 50 mL culture volume. Unless otherwise stated, cultures were incubated at 37°C, 45°C, or 50°C in a closed, dark incubator using the New Brunswick Innova 40 incubator. Conidia were harvested from 7-day-old plates by gentle flooding with sterile 0.01% (v/v) Tween-80, filtered through sterile Miracloth to remove hyphal fragments, counted using a hemocytometer, and used immediately or stored at -80°C until use. All other strains and plasmids generated from the study are listed in Table 2.

**Table 2.** List of strains and plasmids used for this study. For each row, the first column lists the strain or plasmid, the second column provides its description, and the third column indicates the corresponding reference or source for strain construction and plasmid generation.

| Strains | Description | References / sources |
| --- | --- | --- |
| *Escherichia coli* DH5α | F- Φ80lacZΔM15 Δ(lacZYA-argF) U169 recA1 endA1 hsdR17 | Invitrogen |
| *Agrobacterium tumefaciens* LBA4404 | Ach5 with pAch5, Δtra Δocc ΔT<sub>L</sub> ΔT<sub>R</sub> Rif<sup>r</sup> | (57) |
| *Thermothelomyces thermophilus* ATCC 42464 | Wild-type reference strain used as the parental background in this study | (45) |
| MJK20.2 (Δku80) | Mutant strain of *T. thermophilus* having deletion of *ku80* gene (MYCTH_2118116) | (45) |
| Δech | Mutant strain of *T. thermophilus* having deletion of *ech* (MYCTH_2303755) and *ku80* (MYCTH_2118116) genes | This study |
| Δmfs | Mutant strain of *T. thermophilus* having deletion of *mfs* (MYCTH_116881) and *ku80* (MYCTH_2118116) genes | This study |
| Δnrc-1 | Mutant strain of *T. thermophilus* having deletion of *nrc-1* (MYCTH_2305107) and *ku80* (MYCTH_2118116) genes | This study |
| Δpty-5 | Mutant strain of *T. thermophilus* having deletion of *pty-5* (MYCTH_2300403) and *ku80* (MYCTH_2118116) genes | This study |
| Δsyg1 | Mutant strain of *T. thermophilus* having deletion of *syg1* (MYCTH_100809) and *ku80* (MYCTH_2118116) genes | This study |
| ΔMYCTH_2313252 | Mutant strain of *T. thermophilus* having deletion of *MYCTH_2313252* and *ku80* (MYCTH_2118116) genes | This study |
| ΔzincTF | Mutant strain of *T. thermophilus* having deletion of *zincTF* (MYCTH_2139072) and *ku80* (MYCTH_2118116) genes | This study |
| Δzip | Mutant strain of *T. thermophilus* having deletion of *zip* (ΔMYCTH_2307966) and *ku80* (MYCTH_2118116) genes | This study |
| Plasmids | Description | References / sources |
| pCambia 1302 | Kanamycin and hygromycin resistant vector containing EGFP | (58) |
| pLH159RB | Plasmids for *A. tumefaciens*-mediated | (19) |
|  | transformation with barcoded T-DNA |  |
| pOORB007 | pCAMBIA1302-based plasmid bearing the *nrc-1* deletion cassette, consisting of ~1 kb upstream and downstream homology arms flanking the hph selectable marker in place of the *nrc-1* ORF | This study |
| pOORB008 | pCAMBIA1302-based plasmid bearing the *syg1* deletion cassette, consisting of ~1 kb upstream and downstream homology arms flanking the hph selectable marker in place of the *syg1* ORF | This study |
| pOORB009 | pCAMBIA1302-based plasmid bearing the *ech* deletion cassette, consisting of ~1 kb upstream and downstream homology arms flanking the hph selectable marker in place of the *ech* ORF | This study |
| pOORB012 | pCAMBIA1302-based plasmid bearing the *zincTF* deletion cassette, consisting of ~1 kb upstream and downstream homology arms flanking the hph selectable marker in place of the *zincTF* ORF | This study |
| pOORB014 | pCAMBIA1302-based plasmid bearing the *pty-5* deletion cassette, consisting of ~1 kb upstream and downstream homology arms flanking the hph selectable marker in place of the *pty-5* ORF | This study |
| pOORB018 | pCAMBIA1302-based plasmid bearing the *mfs* deletion cassette, consisting of ~1 kb upstream and downstream homology arms flanking the hph selectable marker in place of the *mfs* ORF | This study |
| pOORB020 | pCAMBIA1302-based plasmid bearing the MYCTH_2313252 deletion cassette, consisting of ~1 kb upstream and downstream homology arms flanking the hph selectable marker in place of the MYCTH_2313252 ORF | This study |
| pOORB021 | pCAMBIA1302-based plasmid bearing the *zip* deletion cassette, consisting of ~1 kb upstream and downstream homology arms flanking the hph selectable marker in place of the *zip* ORF | This study |

### Temperature-dependent growth and development assays of wild-type *T. thermophilus*

To characterize thermal performance, conidia were inoculated onto solid VMM + 2% sucrose plates at a final density of 1.2×10^6^ conidia/spot and incubated at 37°C, 45°C, or 50°C for 168 h in a closed incubator under standard dark conditions. Growth was observed daily using a flatbed scanner. Colony pigmentation and aerial hyphae formation were documented under standardized lighting conditions. Conidiation output was quantified from cultures grown on solid medium for 168 h. Conidia were harvested as described above and counted using a hemocytometer in triplicate biological replicates per temperature. For liquid growth assays, conidia were inoculated at 1×10⁶ conidia/mL into 50 mL VMM + 2% sucrose in 250 mL Erlenmeyer flasks and incubated at 30°C, 37°C, 45°C, 50°C, or 55°C for 24 h at 200 rpm under standard dark conditions. Biomass accumulation was quantified by dry weight after filtration onto pre-weighed Whatman GF/A filters and dried at 80°C until a constant mass was achieved.

### Construction and Tnseq of the insertional mutant pool

We generated genome-wide random insertional mutant libraries in *Thermothelomyces thermophilus* ATCC 42464 using *Agrobacterium tumefaciens*–mediated transformation. To obtain fresh starting material, we prepared conidia on VMM containing 2% sucrose and 2% agar and incubated the plates at 37°C for 5–7 days in the dark. Conidia were harvested in sterile water, filtered through Miracloth to remove hyphal debris, counted with a hemocytometer, and adjusted to ∼4×10^7^ conidia/mL. For *Agrobacteriu*m transformation, we used *the A. tumefaciens* strain EHA105, which carries a binary vector encoding a barcoded T-DNA cassette and a hygromycin resistance marker (19). The strain was cultured in low-salt LB with kanamycin in an exponential phase (OD_600_ ∼0.5–0.8), then transferred to induction medium (IM) containing 19.5 g/L 2-(N-morpholino)ethane sulfonic acid (MES) at pH 5.5 supplemented with 500 μM acetosyringone for 18 h to induce virulence following the modified method of Michielse *et. al.* (48). Grown *A. tumefaciens* (OD_600_ ∼0.3–0.7) was mixed with *T. thermophilus* conidia at a 1:1.5 ratio and co-incubated for 1 h at room temperature with gentle mixing to prevent settling.

We spread 1 mL of the co-culture mixture onto the surface of a sterile ultra clear cellophane (#RPI-1090) placed on a 150 × 15 mm IM agar plate supplemented with 500 μM acetosyringone. Plates were incubated at 28°C for 96 h before transferring the cellophane overlays to VMM + 2% sucrose selection plates containing 200 μg/mL hygromycin and 100 μg/mL carbenicillin. Plates were incubated at 37°C until hygromycin-resistant colonies emerged (4–7 days), and conidiation was allowed to proceed for an additional ∼14 days. This full workflow was performed twice, generating two independent mutant pools: Batch 1 (∼4,000 transformants) and Batch 2 (∼13,000 transformants). For each batch, conidia from all selection plates were harvested, pooled, filtered through Miracloth, and stored as 10% glycerol stocks at –80°C. Together, these two batches comprised ∼20,000 unique insertion mutants.

For mapping T-DNA insertion sites (Tnseq), for each batch we inoculated pooled mutant conidia (1×10⁶ conidia/mL) into 100 mL VMM + 2% sucrose and grew the culture for 24 h at 37°C and 200 rpm in the dark before extracting genomic DNA using the Quick-DNA Fungal/Bacterial Miniprep Kit (Zymo Research). No dropout mix was used at any stage, as *T. thermophilus* is not maintained as an auxotrophic system; transformants were selected solely on hygromycin-containing medium, and only disruptions in essential genes would be expected to drop out of the pooled library. We verified DNA integrity by agarose gel electrophoresis and quantified with a Qubit dsDNA HS assay. We prepared T-DNA genome junction sequencing libraries following the general TnSeq workflow of Wetmore et al. (2), but using an enzymatic-fragmentation workflow based on the NEBNext Ultra II FS DNA Library Prep Kit with Sample Purification Beads for Illumina (New England Biolabs #E6177S). Library preparation followed the manufacturer’s instructions with modifications appropriate for TnSeq-style T-DNA junction enrichment as described previously (49). Briefly, FS-based fragmentation and end-repair, adaptor ligation, and dual-SPRI size selection were performed using components of the NEBNext kit, after which nested PCR enrichment of junction fragments was carried out using the Nspacer_barseq_universal and P7_MOD_TS_index primers (sequences in Supplementary File 4). Final libraries were sequenced as paired-end 150 bp reads on an Illumina MiSeq at the UC Berkeley Vincent J. Coates Genomics Sequencing Laboratory.

### Tnseq analysis/mapping of the insertional mutant pool

To determine the genomic coordinates of each barcoded insertion, sequencing reads from Tnseq from Batch 1 and Batch 2 (see above) were combined and then processed using a modified version of the pipeline described by Kim et al. (50). All custom modifications and scripts used in this study are available in the accompanying GitHub repository (https://github.com/oaogunyewo-git/Thermothelomyces-Tnseq-project/tree/main/TnSeq_BarSeq_sequencings_notebooks_and_scripts) and permanently archived on Zenodo (51). Briefly, the algorithm first identified reads containing the T-DNA right-arm junction sequence, allowing up to two mismatches to accommodate sequencing error, and extracted the adjacent genomic flanking sequence. These flanking sequences were aligned to the *T. thermophilus* reference genome (assembly MycTherm_2019, available at the DOE JGI MycoCosm portal) (13) using BLASTn with a minimum alignment identity of 95% and an E-value threshold of 0.1. A barcode was considered to map uniquely only if the highest-scoring genome alignment exceeded the second-best alignment by at least 10 bitscore units. Barcodes differing by single-base sequencing errors were collapsed, and barcodes mapping to nearly identical positions (<10 bp apart) were resolved by retaining the most abundantly supported locus. Barcodes classified as multi-locus, ambiguous, or insert-derived were excluded. Only single-locus barcodes supported by at least one uniquely mapped read were retained. Each insertion was assigned a classification (single, multi-locus, concatameric, or ambiguous) based on mapping structure, and only single-locus insertions were retained for downstream mutant fitness calculations. The final poolfile reported for each barcode, the chromosomal coordinate, strand orientation, gene annotation, and corresponding read abundance. The resulting poolfile was then annotated by intersecting insertion coordinates with gene boundaries defined in the *T. thermophilus* genome annotation (GFF) file, thereby assigning each barcode to a specific gene (or intergenic region) and producing a final gene-level insertion index used for subsequent fitness scoring. Repetitive elements in the *T. thermophilus* reference genome were identified using RepeatMasker v4.2.2 (52) with the Dfam repeat database (53) under default parameters.

### High-throughput phenotyping and quantification of temperature-dependent fitness in the insertional mutant pool

For a given replicate of quantification of abundance of mutant across the pool of insertional mutants (Barseq), we proceeded as follows. A 1 mL frozen aliquot of a given mutant pool (Batch 1 or Batch 2; see above) was spread onto two 150 x 15 mm VMM +2% sucrose agar plates (500 uL per plate) and incubated at 37°C for 6 days in the dark to allow full conidiation. Conidia harvested from these expanded plate cultures were pooled and split into two inocula for growth: 50 mL liquid VMM + 2% sucrose was inoculated with 1×10⁶ conidia/mL in each case, after which one was incubated at 37°C and the other at 50°C for 24 h at 200 rpm under dark conditions. For each, mycelial biomass was harvested for genomic DNA extraction using Quick-DNA Fungal/Bacterial Miniprep Kit (Zymo Research D6005). Unique barcode regions were amplified using Q5 High-Fidelity DNA Polymerase with GC Enhancer (NEB M0491S) and primers targeting constant priming sites flanking each barcode, yielding a ∼185 bp Illumina-ready amplicon (54). Amplicons were purified using Zymo DNA Clean & Concentrator (D4014), quantified using a Qubit 3.0 fluorometer, pooled in equimolar ratios within each batch. Replicates of Batch 1 (the ∼4,000-mutant library) were sequenced on an Illumina MiSeq platform, and replicates of Batch 2 (the ∼13,000-mutant library) were sequenced on an Illumina NextSeq P2 platform (100-cycle single-end). When necessary, libraries were size-selected using a Pippin Prep (Sage Biosciences) and spiked with 15% PhiX to increase base diversity.

Sequencing reads were processed using a modified RBseq_Count_BarCodes pipeline (51). Barcode counts were normalized by total-count scaling and linked to genomic insertion sites via the annotated poolfile; gene-level abundance estimates were derived from uniquely mapped, single-locus barcodes only. As an internal quality control, genome-wide insertion density was summarized in 50-kb chromosomal bins, and replicate concordance was assessed by pairwise scatter plotting of normalized barcode abundances with coefficients of determination (R²) from ordinary least-squares regression. Barcodes whose insertion position from Tnseq mapped to multiple promoters or genes were eliminated from downstream analysis, as was any gene for which all insertion mutants in all replicates had fewer than 5 observed reads.

To test for differential abundance between temperatures of the mutants of a given gene, we proceeded as follows. We calculated the mean normalized abundance across all insertional mutants from all 37°C replicates and divided the value of the normalized abundance from each insertional mutant in each 50°C replicate by this value. Log₂ transformation of these ratios yielded per-gene fold-change estimates. For this baseline-normalized analysis, statistical testing was restricted to genes represented by ≤15 uniquely mapped barcodes, to prevent genes with very high insertion density from disproportionately influencing variance structure. For each such gene, a one-sample Wilcoxon signed-rank test was used to determine whether the median log₂(50°C/37°C) differed from 0 (i.e., ratio = 1). P-values were corrected using the Benjamini–Hochberg procedure, and genes were designated as temperature-responsive when the absolute value of the mean log₂(50°C/37°C) ≥ 1 and FDR < 0.05. We also computed the standard deviation and coefficient of variation (CV%) across all available replicate log₂(50°C/37°C) measurements.

### Single-gene validation of screen hits

#### Construction of single-gene deletion mutants

Single-gene knockout mutants were generated in the Δ*ku80* background of *T. thermophilus* to promote homologous recombination (55). For each target gene, we PCR-amplified the upstream and downstream genomic flanking regions, typically 500 bp to 1 kb each, depending on the locus. We assembled these flanking regions on either side of the hygromycin resistance marker (hph), which was amplified from the pLH159RB plasmid. The flanking regions and hph cassette were inserted into the pCAMBIA1302 binary vector, which had been linearized by *Xho*I and *BssH*II digestion to remove the native GFP cassette and create compatible ends for assembly. Constructs were assembled using NEBuilder HiFi DNA Assembly Master Mix (New England Biolabs, #E5520S). The assembled plasmids were transformed into *Escherichia coli* DH5α, and transformants were selected on LB agar supplemented with 50 µg/mL kanamycin. We screened positive colonies by colony PCR, and plasmid DNA was purified using a miniprep kit (QIAprep Spin Miniprep Kit #27106). Correct assembly was verified by diagnostic restriction enzyme digest and Sanger sequencing across the junction regions to confirm integrity of the flanking regions and hph cassette. The complete list of plasmids generated in this study is provided in Table 2, and all primers used for cassette construction are listed in Supplementary File 5.

The verified pCAMBIA1302-based deletion vectors were introduced into *Agrobacterium tumefaciens* strain LBA4404 by electroporation. Transformants were recovered on low-salt LB agar supplemented with 100 μg/mL kanamycin and 50 μg/mL rifampicin to select for plasmid maintenance and the bacterial background, respectively. Gene deletion in *T. thermophilus* was then carried out by *Agrobacterium*-mediated transformation following the protocol of Michielse *et al.,* 2008 (48). Briefly, freshly harvested conidia (∼4×10⁷ per plate) were co-cultivated with *A. tumefaciens* on cellophane membrane placed directly on IM agar plates containing 500 μM acetosyringone and incubated at 28°C for 96 h as above. After a 96 h incubation period, the cellophane membrane containing the grown co-cultivation mixture was subsequently transferred to the selection plates (VMM+ 2% sucrose supplemented with 200 μg/mL hygromycin and 100 μg/mL carbenicillin) and incubated at 37°C for 4–5 days to recover primary transformants. Hygromycin-resistant colonies were then re-streaked to secondary selection plates containing the same antibiotic and incubated for an additional 10 days to promote full conidial development and ensure stable genomic integration prior to strain purification and downstream verification.

#### Molecular verification of single-gene validation mutants

To verify each gene deletion, we first grew conidiospores from the hygromycin-resistant transformants in 50 mL VMM + 2% sucrose for 72 h at 37°C and 200 rpm in the dark and extracted genomic DNA using the Quick-DNA Fungal/Bacterial Miniprep Kit (Zymo Research, D6005). We confirmed the gene deletions using a two-step PCR strategy. First, we performed 5′ and 3′ verification PCRs using primer pairs located outside the upstream and downstream flanking regions of the targeted locus. This allowed us to distinguish between the wild-type allele and the deletion allele based on product size. For each mutant, we observed a single band corresponding to the expected size of the deletion allele, which differed from the native allele band amplified from the Δ*ku80* parental strain. Second, we used internal ORF-specific primers to test for the presence or absence of the native gene. The deletion strains showed no internal ORF product, while the Δku80 parental strain produced the expected amplicon, confirming loss of the target coding region. PCRs were performed using Q5 High-Fidelity DNA Polymerase (NEB, M0491S) with GC enhancer, and products were resolved on 1% agarose gels. Strains that met both criteria—correct external PCR size and absence of the internal ORF—were purified further by single-conidium isolation to ensure clonal homogeneity. For a subset of mutants, we also sequenced the cassette–genome junctions to confirm precise integration and rule out partial or ectopic events. Primer sequences used for verification are provided in Supplementary File 5.

#### Phenotypic validation of single-gene validation mutants

For the liquid growth assay of single-gene validation mutants, because strains differed in germination efficiency at 37°C and 50°C, we designed a scheme in which all strains began the hyphal-growth phase with a similar biomass input, allowing us to assess post-germination hyphal growth independently of initial germination rate. We achieved this by pre-germinating each strain and replicate in turn at 45°C and ensuring consistent growth-ready inocula in liquid culture, as follows. For a given replicate, conidia were first harvested from fully sporulated cultures, washed once in sterile water, and counted using a hemocytometer. We next used this to establish a pre-culture by inoculating 10 mL of the conidial suspension (∼2 x 10^7^ condiospores/mL) into 90 mL of VMM + 2% sucrose in a 500 mL Erlenmeyer flask, sealing with foil, and pre-germinating for 24 h at 45°C with shaking at 200 rpm under standard dark conditions. We then dispensed 5 mL pre-germinated culture (containing mycelium suspended in medium) into each of three 250 mL Erlenmeyer flasks containing 45 mL fresh VMM + 2% sucrose to establish technical replicate cultures. Replicate flasks were incubated in VMM + 2% sucrose at 37°C or 50°C for 18 h with shaking at 200 rpm under dark conditions. At the end of the incubation, mycelia were harvested by filtration through Miracloth, dried overnight at 50°C, and weighed to obtain final dry biomass (mg). Following incubation, mycelia were collected by filtration through Miracloth, washed with sterile water, and dried overnight at 50°C to constant weight. The dry biomass (mg) was recorded for each culture. In a separate checking step for each replicate, we also took an additional 5 mL of the pre-germinated culture (apart from the inocula used for growth assays), dried it overnight at 50°C, and weighed it. The results from these checks revealed little variation between replicates or genotypes (7.0 – 11.0 mg), confirming that 5-mL aliquots used as inocula for growth would be consistent with respect to growth-ready biomass. Biological replicates were performed with *n* ≥ 3 per strain per condition. For visualization in heatmaps, replicate measurements were aggregated as mean values. Full replicate-level data are provided in Supplementary File 4.

Germination efficiency was assessed as follows. For a given replicate, 150 µL of a conidial suspension (∼2 x 10^7^ condiospores/mL) was inoculated into 1.5 mL VMM + 2% sucrose in each well of a 24-well plate (Cellvis #P24-1.5H-N). Plates were incubated at 37°C or 50°C with shaking at 200 rpm for 10 h in the dark. To ensure uniform sampling, wells were briefly mixed by pipetting with a P1000 immediately before microscopy. This mixing step resuspended settled conidia and did not alter germination conditions. For microscopy, we withdrew 10 μL of the culture from each well and mounted it on a microscope slide for imaging. Germination was scored using a compound light microscope (National Ecoline D-ELDB Binocular Digital Microscope) at 400× magnification under brightfield illumination. A conidium was defined as germinated when the emerging germ tube exceeded the diameter of the spore body. For each replicate, ≥200 conidia were counted manually to calculate percent germination. Each strain and temperature condition was assayed in ≥ 3 biological replicates.

### Assays of zinc effects on germination

Zinc-dependent effects on germination were assessed in liquid VMM supplemented with 2% sucrose using *T. thermophilus* Δ*ku*80 or Δ*zip* strains, as indicated. Vogel’s minimal medium was prepared with graded ZnSO₄ concentrations by adjusting the trace element stock to yield final zinc levels of 0.5, 1, 3, 5, or 10 mg/L ZnSO₄ in the working 1× medium. (Standard VMM contains 5 mg/L ZnSO_4_.) For each zinc condition, 150 µL of a conidial suspension (4 x 10^7^ condiospores/mL) was inoculated into 1.5 mL zinc-adjusted Vogel’s medium in each well of a 24-well plate (Cellvis #P24-1.5H-N). Plates were incubated at 37°C or 50°C with shaking at 200 rpm for 10 h in the dark. Wells were briefly mixed by pipetting with a P1000 immediately prior to sampling to ensure uniform suspension. For microscopy, 10 µL of culture was mounted on a glass slide and imaged using a compound light microscope at 400× magnification under brightfield illumination. A conidium was considered germinated when the germ tube length exceeded the diameter of the spore body. For each replicate, at least 200 conidia were manually counted to calculate the percent germination. Each zinc condition and temperature combination was assayed in ≥3 biological replicates. For Δ*zip* assays shown in Figure 5B, germination was quantified at low (0.5 mg/L) and high (10 mg/L) zinc concentrations using the same conditions, incubation times, and scoring criteria as for wild-type assays.

### Statistical analysis

All data processing and statistical analyses were performed in Python (v3.10) using the pandas, numpy, scipy, statsmodels, matplotlib, and seaborn packages available in the scientific Python ecosystem (56). Barcode abundance tables from the pooled mutant screens were normalized by total-count scaling. Gene-level fitness effects were quantified using two complementary statistical frameworks: (i) a Mann–Whitney U test comparing barcode abundance distributions between 37°C and 50°C, and (ii) a baseline-normalized log₂ fold-change approach in which the mean normalized abundance across 37°C replicates was used as a reference, and a one-sample Wilcoxon signed-rank test assessed whether the median log₂(50°C/37°C) differed from zero. Resulting p-values were corrected for multiple testing using Benjamini–Hochberg false discovery rate (FDR) control, and genes were classified as temperature-responsive when mean log₂FC ≥ 1 and FDR < 0.05. Biological replicate reproducibility was evaluated by pairwise scatter analysis of normalized barcode abundances, with coefficients of determination (R²) computed from ordinary least-squares regression. Volcano plots and coefficient-of-variation (CV) summaries were generated directly from the baseline-normalized log₂ fold-change tables.

For germination and growth phenotyping assays, temperature and genotype effects were assessed using two-way ANOVA (Strain × Temperature) implemented in the statsmodels package within the Python scientific ecosystem (56). Where a significant strain × temperature interaction was detected, post-hoc comparisons were conducted. For growth, Tukey’s HSD was applied to the strain factor to identify genotypes that differed from Δ*ku80* across temperatures, with significance determined by the Tukey-adjusted p-value. Heatmaps were generated with seaborn, and significance markers were placed directly on the row margins. All figures were generated in Python and finalized in Inkscape software version 1.4.2.

## Data availability

Raw sequencing data are available from the NCBI Sequence Read Archive (SRA) under accession number PRJNA1355313. Germination assay micrographs are available on Figshare at https://doi.org/10.6084/m9.figshare.30593345. T-DNA sequencing data are in Supplementary File 1, and barcode sequencing data are in Supplementary File 3. Growth, conidiation and germination data for wild-type and single-gene validation mutants are in Supplementary File 4. Plasmids generated in this study are listed in Table 2, and all primers used for cassette construction and strain verification are provided in Supplementary File 5.

## Supporting information

All supplementary files

## Acknowledgements

We thank Prof. Louise Glass, Dr. Florian Drescher, Dr. José Manuel Villalobos-Escobedo, and Dr. Adam Deutschbauer for invaluable guidance and support of this work. We also thank the members of the Brem Lab for insightful suggestions. O.A.O. acknowledges financial support from the Energy & Biosciences Institute (EBI) at UC Berkeley through the EBI-Shell Postdoctoral Fellow program. This work was supported in part by NIH R01GM120430 to R.B.B.

## Supplementary figure captions

**Supplementary Figure 1. Reproducibility of barcode abundance measurements across biological replicates.** Data are as in Figure 3B of the main text, except that panels A–C correspond to replicate comparisons at 37°C for the Batch 1 mutant pool (sets A1–A2, B1–B2, and C1–C2); D–E show replicate comparisons at 50°C for the Batch 1 mutant pool (sets A1–A2 and B1–B2); F-G correspond to replicate comparisons at 37°C for the Batch 2 mutant pool; H-I correspond to replicate comparisons at 50°C for the Batch 2 mutant pool. The coefficient of determination (R²) for each comparison is shown in the top left of each plot.

**Supplementary Figure 2. PCR confirmation of targeted gene deletions in *T. thermophilus*.** Diagnostic PCR was used to verify homologous replacement of each target locus in the deletion strains (Δ*mfs*, Δ*syg1*, Δ*nrc-1*, Δ*pty-5*, Δ*zincTF*, Δ*zip*, *ΔMYCTH_2313252* and Δ*ech*). For each gene, lanes labeled C correspond to the parental Δ*ku80* control, and adjacent lanes show amplicons from independent deletion-mutant isolates. Loss of the wild-type band and/or the presence of the expected deletion-specific product confirms successful gene replacement at the target locus. M refers to the DNA size marker. The expected size bands for each of the wild type and transformant PCRs are listed in Supplementary File 4.

## Supplementary file captions

**Supplementary File S1. T-DNA junction sequencing of *T. thermophilus* insertional mutants.** Listed are all mapped insertion mutants in the *T. thermophilus* genome. Columns report barcode and reverse-complement sequences (barcode, rcbarcode), total read support (nTot) and primary-locus support (n), genomic location (scaffold, strand, pos), mapping classification (type), read support for the main mapped site (nMainLocation) and for T-DNA border–derived reads (nInsert), all alternate genomic alignments (All genomic mappings), and annotated overlapping or nearest gene (gene).

**Supplementary File S2. Genome-wide RepeatMasker annotations for the *T. thermophilus* reference genome.** This file contains RepeatMasker v4.2.2 annotations generated using the Dfam repeat database. Columns report chromosome, start coordinate, end coordinate, repeat feature classification, motif annotation, score, strand, and full RepeatMasker attribute string.

**Supplementary File S3. Genome-wide BarSeq fitness profiles of *T. thermophilus* at 37^°^C and 50^°^C.** This file contains three worksheets reporting results of gene-level tests for differential abundance between temperatures from BarSeq data. In each sheet, GeneID is the annotated gene identifier; Mean_log2FC is the mean log₂ fold change in mutant abundance (50°C/37°C); P-Value is the nominal *p*-value from the Wilcoxon rank-sum test for differential abundance between temperatures; Adjusted_P-Value is the multiple-testing–corrected *p*-value (Benjamini–Hochberg); protein_names is the predicted or annotated protein name; Std_log2FC is the standard deviation of mean log₂ fold change in mutant abundance (50°C/37°C) across mutants and replicates; and CV_log2FC_percent is the coefficient of variation of mean log₂ fold change in mutant abundance (50°C/37°C) across mutants and replicates, expressed as a percentage. Worksheet 1 includes results from all genes with BarSeq data that met our quality control threshold (≤15 detected insertions; see Methods). Worksheet 2 reports genes whose mutants exhibited temperature-responsive fitness that met significance and effect-size thresholds for which mean log₂ fold change in mutant abundance (50°C/37°C) ≥ 1. Worksheet 3 reports genes that met significance and effect-size thresholds for which mean log₂ fold change in mutant abundance (50°C/37°C) ≤ –1.

**Supplementary File S4. Raw measurements of conidiation, biomass accumulation, and germination of wild-type and single-gene validation mutant *T. thermophilus*.** This file contains five worksheets reporting unprocessed measurements underlying Figures 1, 4, and 5. Worksheet 1 reports raw counts of conidia harvested from wild-type cultures grown on solid VMM + 2% sucrose at 37°C, 45°C, and 50°C. Columns report temperature and conidia count. Worksheet 2 reports dry-weight biomass measurements for wild-type cultures grown in liquid VMM containing 2% sucrose at 30°C, 37°C, 45°C, 50°C, and 55°C. Columns report temperature and growth (mg). Worksheet 3 reports Biomass accumulation dry-weight biomass measurements for all deletion mutants and the Δ*ku*80 parent grown in liquid VMM + 2% sucrose at 37°C and 50°C. Columns report strain, temperature, and growth (mg). Worksheet 4 reports microscopy-based germination measurements for all deletion mutants and the Δku80 parent at 37°C and 50°C. Columns report strain, temperature, total number of conidia, number of germinated conidia, and percentage germination. Worksheet 5 reports the raw germination data for Δku80 conidia exposed to varying Zn concentrations at 37°C and 50°C. Columns report zinc concentration, temperature, total number of conidia, number of germinated conidia, and percentage germination.

**Supplementary File S5. Primer sequences used for construction of deletion cassettes and validation of gene deletions by PCR.** This file contains three worksheets detailing the oligonucleotides used in this study for TnSeq/BarSeq library preparation, deletion-cassette construction, and PCR-based mutant verification. Worksheet 1 reports the primers used for Tnseq and Barseq amplifications, indexing, and nested enrichment of T-DNA junction fragments. The columns report primer_name, primer_sequence, index_sequence (for multiplexing), and index_reverse (custom P1 index primer). Worksheet 2 reports the primers used to assemble deletion cassettes and to verify correct integration in transformants by PCR. Columns report GeneID, primer_name, primer_sequence, and purpose (e.g., 5′ flank amplification, junction PCR, confirmation PCR). Worksheet 3 reports the expected amplicon sizes for wild-type (Δku80 background) and deletion mutants after cassette integration. Columns report gene, primer_set, expected_size_in_Δku80 (kb), and expected_size_in_mutant (kb).

## Notes

### Competing Interest Statement

The authors have declared no competing interest.

### Summary of Updates

We investigated the origin of a hotspot of Agrobacterium-mediated transposon insertion on Thermothelomyces chromosome 7.

## References

1. Tansey MR, Brock TD. 1972. The Upper Temperature Limit for Eukaryotic Organisms. Proc Natl Acad Sci USA 69:2426.

2. Maheshwari R, Bharadwaj G, Bhat MK. 2000. Thermophilic Fungi: Their Physiology and Enzymes. Microbiol Mol Biol Rev 64:461–488.

3. Wiegant WM. 1992. Growth Characteristics of the Thermophilic Fungus *Scytalidium thermophilum* in Relation to Production of Mushroom Compost. Appl Environ Microbiol 58:1301.

4. Siliakus MF, van der Oost J, Kengen SWM. 2017. Adaptations of archaeal and bacterial membranes to variations in temperature, pH and pressure. Extremophiles 21:651–670.

5. Wang Q, Cen Z, Zhao J. 2015. The survival mechanisms of thermophiles at high temperatures: An angle of omics. Physiology 30: 97–106.

6. Xiao W, Zhang J, Huang J, Xin C, Li MJ, Song Z. 2022. Response and regulatory mechanisms of heat resistance in pathogenic fungi. Appl Microbiol Biotechnol 106:5415.

7. Fabri JHTM, Rocha MC, Fernandes CM, Persinoti GF, Ries LNA, Cunha AF da, Goldman GH, Del Poeta M, Malavazi I. 2021. The Heat Shock Transcription Factor HsfA Is Essential for Thermotolerance and Regulates Cell Wall Integrity in *Aspergillus fumigatus*. Front Microbiol 12:656548.

8. Francisco CS, McDonald BA, Palma-Guerrero J. 2023. A transcription factor and a phosphatase regulate temperature-dependent morphogenesis in the fungal plant pathogen *Zymoseptoria tritici*. Fungal Genet Biol 167:103811.

9. Yaakoub H, Mina S, Calenda A, Bouchara JP, Papon N. 2022. Oxidative stress response pathways in fungi. Cell Mol Life Sci 79:333.

10. Carruthers DN, Lee TS. 2022. Translating advances in microbial bioproduction to sustainable biotechnology. Front Bioeng Biotechnol 10:968437.

11. Adegboye MF, Ojuederie OB, Talia PM, Babalola OO. 2021. Bioprospecting of microbial strains for biofuel production: metabolic engineering, applications, and challenges. Biotechnol Biofuels 14:1–21.

12. Yuan SF, Guo GL, Hwang WS. 2017. Ethanol production from dilute-acid steam-exploded lignocellulosic feedstocks using an isolated multistress-tolerant *Pichia kudriavzevii* strain. Microb Biotechnol 10:1581.

13. Berka RM, Grigoriev I V., Otillar R, Salamov A, Grimwood J, Reid I, Ishmael N, John T, Darmond C, Moisan MC, Henrissat B, Coutinho PM, Lombard V, Natvig DO, Lindquist E, Schmutz J, Lucas S, Harris P, Powlowski J, Bellemare A, Taylor D, Butler G, De Vries RP, Allijn IE, Van Den Brink J, Ushinsky S, Storms R, Powell AJ, Paulsen IT, Elbourne LDH, Baker SE, Magnuson J, Laboissiere S, Clutterbuck AJ, Martinez D, Wogulis M, De Leon AL, Rey MW, Tsang A. 2011. Comparative genomic analysis of the thermophilic biomass-degrading fungi *Myceliophthora thermophila* and *Thielavia terrestris*. Nat Biotechnol 29:922–927.

14. Contato AG, Borelli TC, Buckeridge MS, Rogers J, Hartson S, Prade RA, Polizeli M de LT de M. 2024. Secretome Analysis of Thermothelomyces thermophilus LMBC 162 Cultivated with Tamarindus indica Seeds Reveals CAZymes for Degradation of Lignocellulosic Biomass. J Fungi 10 (121): 1–14.

15. Liu Q, Gao R, Li J, Lin L, Zhao J, Sun W, Tian C. 2017. Development of a genome-editing CRISPR/Cas9 system in thermophilic fungal Myceliophthora species and its application to hyper-cellulase production strain engineering. Biotechnol Biofuels 10:1–14.

16. Liu Q, Zhang Y, Li F, Li J, Sun W, Tian C. 2019. Upgrading of efficient and scalable CRISPR-Cas-mediated technology for genetic engineering in thermophilic fungus *Myceliophthora thermophila*. Biotechnol Biofuels 12:1–19.

17. Zhang C, Li N, Rao L, Li J, Liu Q, Tian C. 2022. Development of an Efficient C-to-T Base-Editing System and Its Application to Cellulase Transcription Factor Precise Engineering in Thermophilic Fungus *Myceliophthora thermophila*. Microbiol Spectr 10:e02321–21.

18. Kwon MJ, Schütze T, Spohner S, Haefner S, Meyer V. 2019. Practical guidance for the implementation of the CRISPR genome editing tool in filamentous fungi. Fungal Biol Biotechnol 6:1–11.

19. Huberman LB, Villalobos-Escobedo JM, Skerker JM, Shi R, Rico-Ramirez AM, Adams C, Arkin AP, Deutschbauer AM, Glass L. 2025. Construction of a randomly barcoded insertional mutant library in the filamentous fungus *Trichoderma atroviride*. bioRxiv 2025.11.30.691285.

20. Xu J, Li J, Lin L, Liu Q, Sun W, Huang B, Tian C. 2015. Development of genetic tools for *Myceliophthora thermophila*. BMC Biotechnol 15:35.

21. Billmyre RB, Craig CJ, Lyon JW, Reichardt C, Kuhn AM, Eickbush MT, Zanders SE. 2025. Landscape of essential growth and fluconazole-resistance genes in the human fungal pathogen *Cryptococcus neoformans*. PLoS Biol 23:e3003184.

22. Segal ES, Gritsenko V, Levitan A, Yadav B, Dror N, Steenwyk JL, Silberberg Y, Mielich K, Rokas A, Gow NAR, Kunze R, Sharan R, Berman J. 2018. Gene essentiality analyzed by in vivo transposon mutagenesis and machine learning in a stable haploid isolate of *candida albicans*. mBio 9:1–21.

23. Michel AH, Hatakeyama R, Kimmig P, Arter M, Peter M, Matos J, De Virgilio C, Kornmann BT. 2017. Functional mapping of yeast genomes by saturated transposition. Elife 6. e23570.

24. Price MN, Wetmore KM, Waters RJ, Callaghan M, Ray J, Liu H, Kuehl J V., Melnyk RA, Lamson JS, Suh Y, Carlson HK, Esquivel Z, Sadeeshkumar H, Chakraborty R, Zane GM, Rubin BE, Wall JD, Visel A, Bristow J, Blow MJ, Arkin AP, Deutschbauer AM. 2018. Mutant phenotypes for thousands of bacterial genes of unknown function. Nature 557:503–509.

25. Blake Billmyre R, Eickbush MT, Craig CJ, Lange JJ, Wood C, Helston RM, Zanders SE. 2022. Genome-wide quantification of contributions to sexual fitness identifies genes required for spore viability and health in fission yeast. PLoS Genet 18:e1010462.

26. Ijadpanahsaravi M, Wösten HAB. 2024. Germination strategies of stress-resistant Aspergillus conidia. Curr Opin Food Sci 57:101169.

27. Wyatt TT, Wösten HAB, Dijksterhuis J. 2013. Fungal spores for dispersion in space and time. Adv Appl Microbiol 85:42–91.

28. Osherov N, May GS. 2001. The molecular mechanisms of conidial germination. FEMS Microbiol Lett 199:153–160.

29. Amir H, Pineau R. 1998. Effects of metals on the germination and growth of fungal isolates from New Caledonian ultramafic soils. Soil Biol Biochem 30:2043–2054.

30. Feigl G, Lehotai N, Molnár Á, Ördög A, Rodríguez-Ruiz M, Palma JM, Corpas FJ, Erdei L, Kolbert Z. 2014. Zinc induces distinct changes in the metabolism of reactive oxygen and nitrogen species (ROS and RNS) in the roots of two Brassica species with different sensitivity to zinc stress. Ann Bot 116:613.

31. Santos TA de O, Soares LW, Oliveira LN, Moraes D, Mendes MS, Soares CM de A, Bailão AM, Bailão MGS. 2024. Zinc Starvation Induces Cell Wall Remodeling and Activates the Antioxidant Defense System in *Fonsecaea pedrosoi*. J Fungi 10:118.

32. Egan MJ, Wang ZY, Jones MA, Smirnoff N, Talbot NJ. 2007. Generation of reactive oxygen species by fungal NADPH oxidases is required for rice blast disease. Proc Natl Acad Sci USA 104:11772–11777.

33. Zhang Z, Chen Y, Li B, Chen T, Tian S. 2020. Reactive oxygen species: A generalist in regulating development and pathogenicity of phytopathogenic fungi. Comput Struct Biotechnol J 18:3344.

34. Fischer MS, Glass NL. 2019. Communicate and Fuse: How Filamentous Fungi Establish and Maintain an Interconnected Mycelial Network. Front Microbiol 10.:619

35. Ghosh A, Servin JA, Park G, Borkovich KA. 2014. Global analysis of serine/threonine and tyrosine protein phosphatase catalytic subunit genes in *Neurospora crassa* reveals interplay between phosphatases and the p38 mitogen-activated protein kinase. G3 (Bethesda) 4:349–365.

36. Desmarini D, Lev S, Furkert D, Crossett B, Saiardi A, Kaufman-Francis K, Li C, Sorrell TC, Wilkinson-White L, Matthews J, Fiedler D, Teresa Djordjevic J. 2020. Ip7-spx domain interaction controls fungal virulence by stabilizing phosphate signaling machinery. mBio 11:1–20.

37. Tamayo E, Gómez-Gallego T, Azcón-Aguilar C, Ferrol N. 2014. Genome-wide analysis of copper, iron and zinc transporters in the arbuscular mycorrhizal fungus *Rhizophagus irregularis*. Front Plant Sci 5:113084.

38. Patkar RN, Ramos-Pamplona M, Gupta AP, Fan Y, Naqvi NI. 2012. Mitochondrial β-oxidation regulates organellar integrity and is necessary for conidial germination and invasive growth in *Magnaporthe oryzae*. Mol Microbiol 86:1345–1363.

39. Michielse CB, Hooykaas PJJ, van den Hondel CAMJJ, Ram AFJ. 2005. Agrobacterium-mediated transformation as a tool for functional genomics in fungi. Curr. Genet. 48:1–17.

40. Gelvin SB. 2003. Agrobacterium-mediated plant transformation: the biology behind the “gene-jockeying” tool. Microbiol Mol Biol Rev 67:16–37.

41. Natasha N, Shahid M, Bibi I, Iqbal J, Khalid S, Murtaza B, Bakhat HF, Farooq ABU, Amjad M, Hammad HM, Niazi NK, Arshad M. 2022. Zinc in soil-plant-human system: A data-analysis review. Sci. Total Environ. 808:152024.

42. Eide DJ. 2006. Zinc transporters and the cellular trafficking of zinc. Biochim Biophys Acta Mol Cell Res 1763:711–722.

43. Wilson S, Bird AJ. 2016. Zinc sensing and regulation in yeast model systems. Arch Biochem Biophys 611:30.

44. Leuenberger P, Ganscha S, Kahraman A, Cappelletti V, Boersema PJ, Von Mering C, Claassen M, Picotti P. 2017. Cell-wide analysis of protein thermal unfolding reveals determinants of thermostability. Science (1979) 355.

45. Drescher F, Li Y, Villalobos-Escobedo JM, Haefner S, Huberman LB, Glass NL. 2024. Transcriptomic and genetic analysis reveals a Zn2Cys6 transcription factor specifically required for conidiation in submerged cultures of *Thermothelomyces thermophilus*. mBio 16(1): e03111–24.

46. Kwon MJ, Schütze T, Spohner S, Haefner S, Meyer V. 2019. Practical guidance for the implementation of the CRISPR genome editing tool in filamentous fungi. Fungal Biol Biotechnol 6:15.

47. Vogel HJ. 1956. A Convenient Growth Medium for *Neurospora crassa*. Microbiol Genet Bull 13:42.

48. Michielse CB, Hooykaas PJJ, van den Hondel CAMJJ, Ram AFJ. 2008. Agrobacterium-mediated transformation of the filamentous fungus *Aspergillus awamori*. Nat Protoc 3:1671–1678.

49. Helmann TC, Filiatrault MJ, Stodghill P V. 2022. Genome-Wide Identification of Genes Important for Growth of *Dickeya dadantii* and *Dickeya dianthicola* in Potato (Solanum tuberosum) Tubers. Front Microbiol 13:778927.

50. Kim J, Coradetti ST, Kim YM, Gao Y, Yaegashi J, Zucker JD, Munoz N, Zink EM, Burnum-Johnson KE, Baker SE, Simmons BA, Skerker JM, Gladden JM, Magnuson JK. 2021. Multi-Omics Driven Metabolic Network Reconstruction and Analysis of Lignocellulosic Carbon Utilization in *Rhodosporidium toruloides*. Front Bioeng Biotechnol 8.

51. oaogunyewo-git. oaogunyewo-git/Thermothelomyces-Tnseq-project: Initial code release for manuscript reproducibility 10.5281/ZENODO.17586924.

52. Flynn JM, Hubley R, Goubert C, Rosen J, Clark AG, Feschotte C, Smit AF. 2020. RepeatModeler2 for automated genomic discovery of transposable element families. Proc Natl Acad Sci USA 117:9451–9457.

53. Hubley R, Finn RD, Clements J, Eddy SR, Jones TA, Bao W, Smit AFA, Wheeler TJ. 2016. The Dfam database of repetitive DNA families. Nucleic Acids Res 44:D81–D89.

54. Coradetti ST, Pinel D, Geiselman GM, Ito M, Mondo SJ, Reilly MC, Cheng Y-F, Bauer S, Grigoriev I V, Gladden JM, Simmons BA, Brem RB, Arkin AP, Skerker JM. 2018. Functional genomics of lipid metabolism in the oleaginous yeast Rhodosporidium toruloides 7:e32110.

55. Siebecker B, Schütze T, Spohner S, Haefner S, Meyer V. 2023. Transcriptomic insights into the roles of the transcription factors Clr1, Clr2 and Clr4 in lignocellulose degradation of the thermophilic fungal platform *Thermothelomyces thermophilus*. Front Bioeng Biotechnol 11: 1279146.

56. Harris CR, Millman KJ, van der Walt SJ, Gommers R, Virtanen P, Cournapeau D, Wieser E, Taylor J, Berg S, Smith NJ, Kern R, Picus M, Hoyer S, van Kerkwijk MH, Brett M, Haldane A, del Río JF, Wiebe M, Peterson P, Gérard-Marchant P, Sheppard K, Reddy T, Weckesser W, Abbasi H, Gohlke C, Oliphant TE. 2020. Array programming with NumPy. Nature 585:357–362.

57. Randhawa A, Ogunyewo OA, Eqbal D, Gupta M, Yazdani SS. 2018. Disruption of zinc finger DNA binding domain in catabolite repressor Mig1 increases growth rate, hyphal branching, and cellulase expression in hypercellulolytic fungus *Penicillium funiculosum* NCIM1228. Biotechnol Biofuels 11:15.

58. Rizwan HM, Yang Q, Yousef AF, Zhang X, Sharif Y, Kaijie J, Shi M, Li H, Munir N, Yang X, Wei X, Oelmüller R, Cheng C, Chen F. 2021. Establishment of a novel and efficient agrobacterium-mediated in planta transformation system for passion fruit (*Passiflora edulis*). Plants 10:2459.

