## Supplementary figures and images for "High-throughput screening reveals mechanisms of environmental control of germination in a fungal thermophile"

### Supplementary Figure S1.pdf

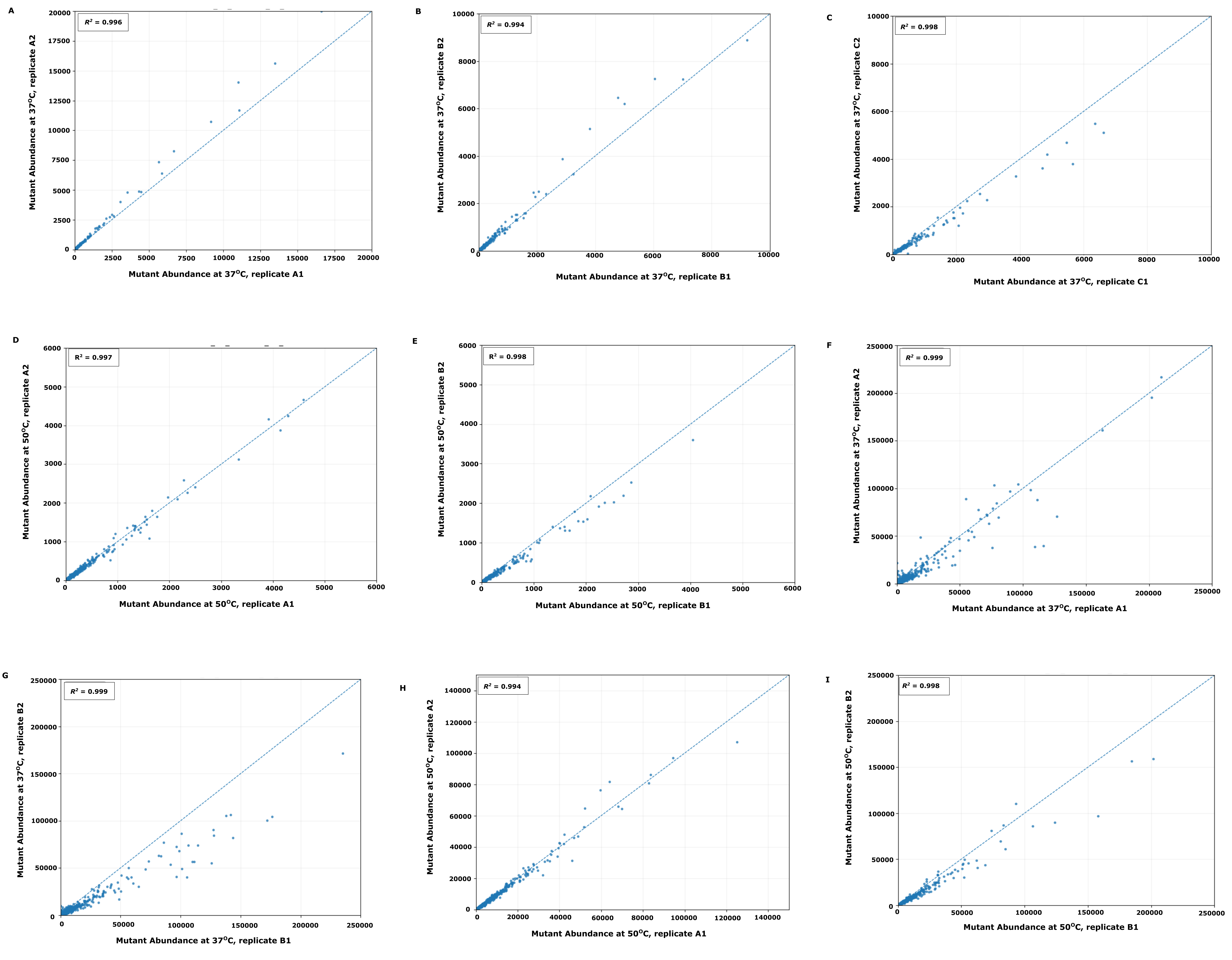

### Supplementary Figure S2.pdf

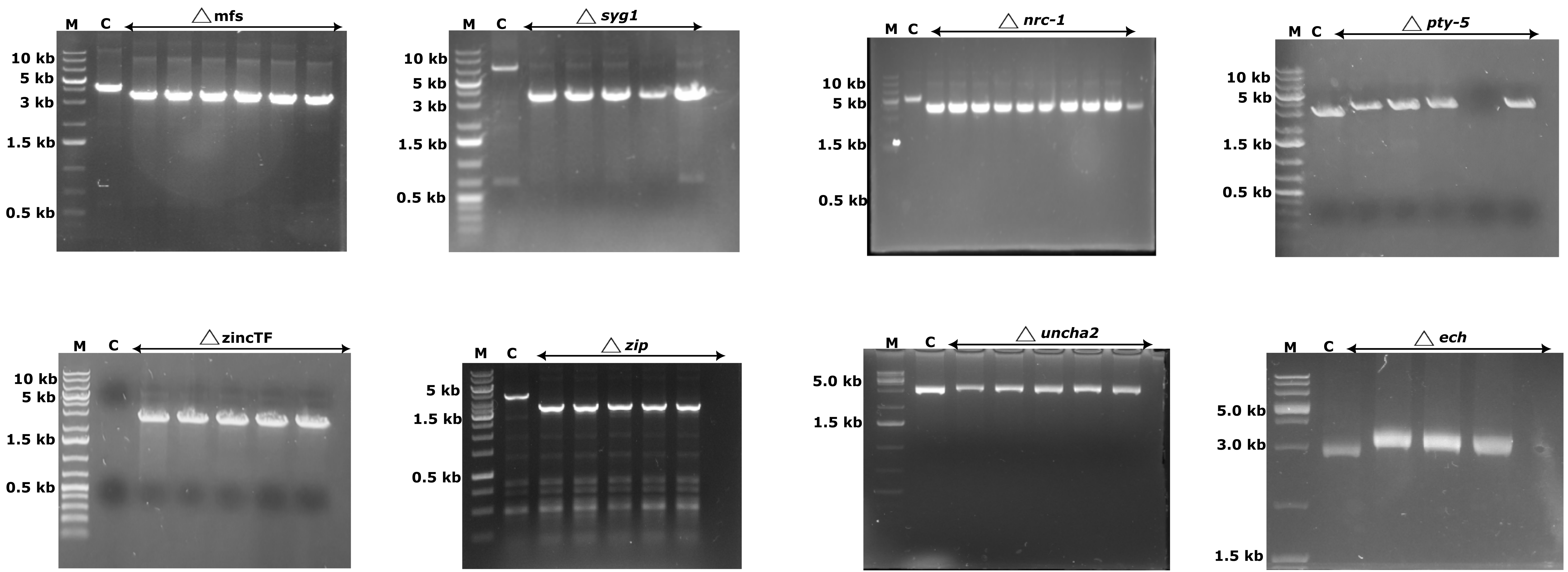
